# The *Stellate* meiotic drive system of *Drosophila melanogaster* is active in contemporary populations

**DOI:** 10.1101/2025.10.02.680149

**Authors:** Benjamin K. McCormick, Daniel A. Barbash, Andrew G. Clark

## Abstract

Meiotic drivers are selfish elements that bias their own transmission so that they are overrepresented among the functional gametes produced. The selective costs imposed by drivers on their hosts may trigger intragenomic conflict, promoting the evolution of suppressors and fueling an ongoing arms race between drivers and suppressors. *Stellate* (*Ste*) is an X-linked tandemly arrayed multicopy gene. Its copy number ranges from 3 to more than 300 among *Drosophila melanogaster* strains from the Global Diversity Lines. In wild-type animals, *Ste* expression is usually suppressed by homologous piRNAs produced from the *Suppressor of Stellate* (*Su(Ste)*) array on the Y chromosome. Derepression of *Ste* in the absence of *Su(Ste)* results in the formation of proteinaceous crystals in spermatocytes, chromatin compaction defects, reductions in fertility, and female-biased sex ratios arising from under-recovery of Y-bearing sperm. Despite extensive study, the function of the *Stellate* array and evolutionary significance of its persistence in the genome have remained elusive. It has been suggested to be a now-inactive relic of an ancient meiotic drive system, as perturbations in lab stocks can produce *Ste*-mediated meiotic distortions. Meiotic drive occurring among natural variants, however, has not been reported. We established crosses between females with high *Ste* copy number X chromosomes and males carrying low *Su(Ste)* copy number Y chromosomes and found that the male progeny displayed non-Mendelian sex chromosome transmission. Importantly, deletion of the *euSte* array in an otherwise matched genetic background rescues this phenotype, demonstrating that *Stellate* is an active driver in contemporary populations.

## Introduction

Meiotic drivers are selfish genetic elements that co-opt gametogenesis to increase their own transmission by ensuring that they are overrepresented among the functional gametes produced^1,2^. This means that, in heterozygotes, the driving allele is inherited more than half the time, while the alternative allele is proportionally undertransmitted. Drivers are thus predicted to rapidly increase in frequency in populations even if they carry substantial costs to host fertility or fitness. Coevolutionary arms race dynamics between the driver and host often promote innovation of drive suppressors through changes in the copy number, sequence, or expression of short RNAs^3–5^ or genes^6–8^. These conflicts are hypothesized to have profound consequences for genome evolution, from reshaping the karyotype^9^ to driving rapid divergence in the sequence of proteins underlying reproduction^10^ and chromosome segregation^11^. Furthermore, disruption of recently-evolved drive suppressors often results in partial or complete sterility^12–14^, leading to the suggestion^e.g. 2^ that drive conflicts may commonly contribute to the emergence and accumulation of hybrid incompatibilities between recently isolated populations and species.

Despite abundant indirect evidence of drive conflicts in many lineages, as well as known drive systems spanning the tree of life, relatively few examples of drivers active in contemporary populations are known. The reasons for this are unclear. Possibilities include biases in detecting drivers, predominance of weak drivers or short-lived conflicts (which resolve through either fixation or suppression), and the context-dependent nature of drive, which may manifest only in some genetic backgrounds or under certain environmental conditions^15–17^. For these reasons, many basic questions about drive remain unanswered: *How common is meiotic drive in natural populations? What features enable a meiotic drive system to persist in a long-term polymorphic state? And what do meiotic drivers look like in natural populations, in terms of strength of drive, allele frequency, stability or resolution of conflict over time, and context dependence?* Answering these questions is essential to understanding this understudied class of selfish elements and their role in shaping the evolution of the genomes that harbor them.

One particularly interesting class of meiotic drivers are sex ratio distorters which, as their name suggests, bias the inheritance of the sex chromosomes^18^. These drivers are not only easier to detect than their autosomal counterparts, and therefore better studied, but they also have distinctive consequences for the populations that carry them. Indeed, a sex ratio distorter that achieves perfect drive (i.e. 100% transmission) is expected to spread rapidly to fixation, ultimately leading to population extinction through the complete loss of one sex. This means that, to avoid this fate and remain active, a sex ratio driver must be limited in some way–whether by constraints inherent to the mechanism of drive, host defenses, or costs to fertility or viability that balance the driver’s transmission advantage.

*Stellate* (*Ste*) is a multicopy protein-coding gene present in two arrays on the X chromosome of *Drosophila melanogaster*, the euchromatic *euSte* array and the pericentric *hetSte* array (Fig. 1A)^19–23^. These genes are specific to this species, having arisen since its split from the *D. simulans* complex ~3 mya^24–26^. *Ste* encodes a 172 aa protein with homology to Ssl, a testis-specific β-subunit of protein kinase CK2^27^. In most situations, expression of *Ste* is repressed by piRNAs generated from homologous Y-linked repeats called *Suppressor of Stellate* (*Su(Ste)*). Derepression of *Ste* in the absence of *Su(Ste)* in males results in meiotic drive disfavoring Y-bearing sperm, the formation of proteinaceous crystals in the developing spermatocytes, as well as reduced fertility and meiotic defects^19,28^. Both *Ste* and *Su(Ste)* exhibit substantial copy number variation both within and between natural populations, with some alleles reaching copy numbers in the hundreds^19,29^. One proposed explanation for this unusual architecture is that *Ste* functions as a sex ratio distorter, with *Ste* and its suppressor locked in an arms race driving progressively higher copy numbers of both elements^30–32^. A model involving a chaseaway process of this sort is supported by the striking observation that, while the primary function of piRNA pathway is thought to be transposon silencing, *Su(Ste)* piRNAs are actually an order of magnitude more abundant than all transposon-directed piRNAs combined^5^.

**Figure 1:**
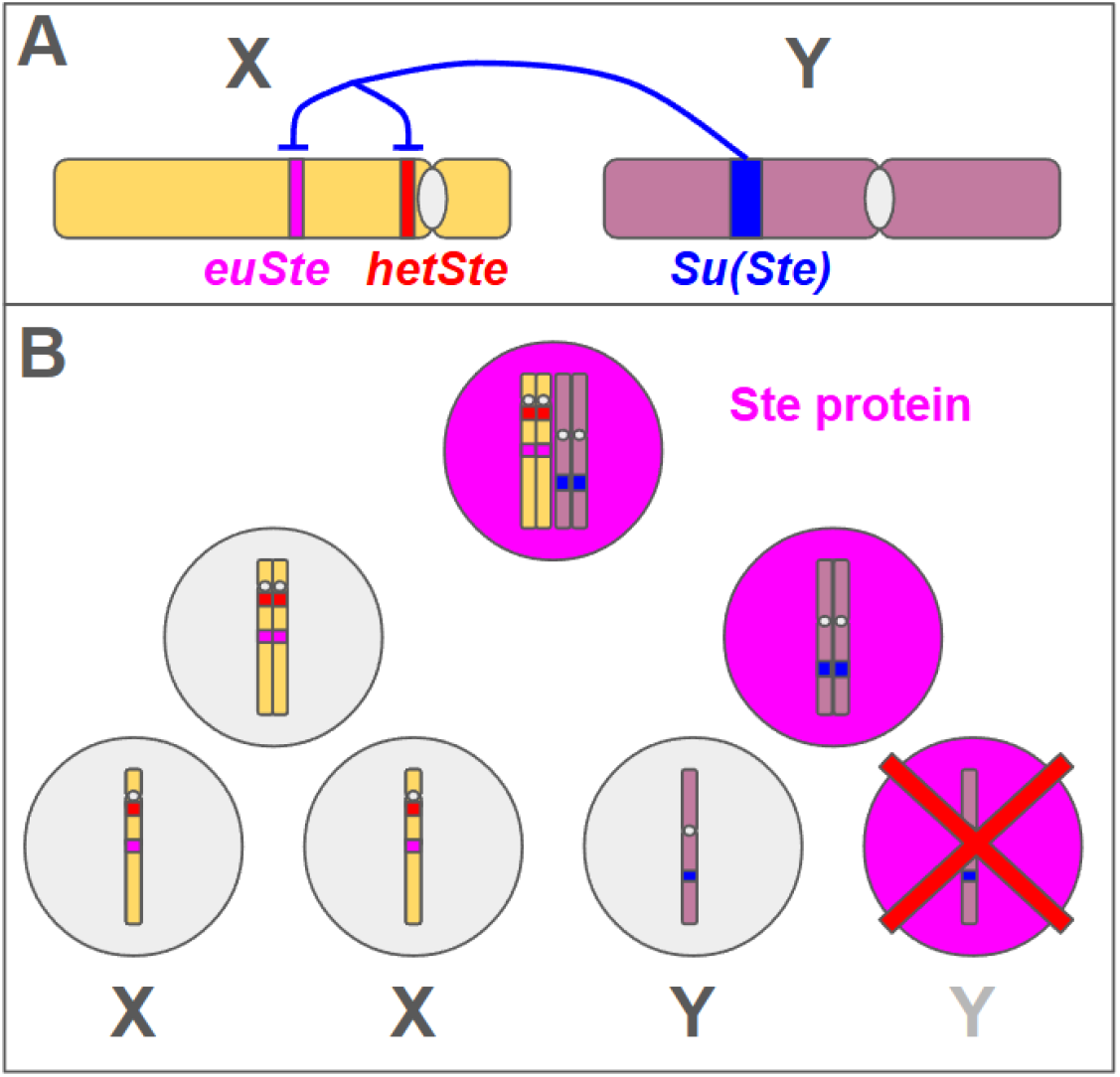
A) Summary of the *Stellate-Suppressor of Stellate* meiotic drive system. The two *Stellate* arrays are found on the X chromosome of *D. melanogaster*; the expression of these genes is generally repressed by the production of piRNAs from the Y-linked *Su(Ste)* array. B) The mechanism of *Ste* drive proposed by Meng & Yamashita (2025). Toxic Ste protein usually segregates asymmetrically during meiosis, being deposited in only one daughter cell per division. In meiosis I, Ste protein is preferentially inherited by the same daughter cell as the Y chromosome. Owing to this bias, the subsequent failure of Ste-bearing spermatids to complete spermiogenesis leads to an under-representation of Y-bearing sperm, producing female-biased sex ratios in the offspring of *Ste*-expressing males.

Furthermore, a recent study^33^ showed that overexpression of *euSte* is necessary and sufficient to produce meiotic drive mediated by preferential cosegregation of Ste protein with the Y chromosome in meiosis I and subsequent killing of the (primarily) Y-bearing daughter cells that inherit it (Fig. 1B). While the longstanding assumption in the literature has been that the *Ste* system is a relic of an ancient, now-suppressed drive conflict^e,g, 1,30,31^, we uncover patterns of copy number variation consistent with active conflict. We further demonstrate that naturally occurring high-copy-number *euSte* alleles produce and are necessary for sex ratio drive in contemporary genetic backgrounds.

## Results

### Patterns of natural variation in the copy number of *Stellate*-family elements in the GDL

To assess population variation of *Ste*-family elements, we estimated their copy number in 48 Global Diversity Lines (GDL)^34^ representing five worldwide populations by two methods: read depth of alignments to a consensus sequence for each element (Supplemental File 1, Table 1) and, for the subset of lines (*n* = 27) for which high quality assemblies were available, by BLAST and phylogenetic clustering (Supplemental File 1, Table 9). In general, these methods produced highly concordant results (Spearman’s rho for *euSte*: 0.91; *hetSte*: 0.80; *Su(Ste)*: 0.59; Supplemental File 1, Table 10), with the discrepancies observed likely arising from assembly errors due to the repetitive structure of the arrays. Disagreement between methods was much more extensive for the Y-linked *Su(Ste)* than for the X-linked *Ste* arrays. This likely results from two factors: 1) the complex structure of the *Su(Ste)* array on the Y chromosome makes it prone to misassembly, and 2) *Su(Ste)* copies exhibit much greater sequence variability than the *Ste* genes, causing copy number estimates based on read depth to systematically undercount highly divergent or degenerate copies, thereby increasing sensitivity to the specific mapping parameters used.

The *euSte* array in the middle of the long arm of the X chromosome exhibits much wider variation in copy number than does the pericentric *hetSte* array (2-293 copies compared to 1-36 copies, respectively; Fig. 2). The open reading frames of these copies are largely intact in both arrays, though pseudogenized copies are substantially more frequent in *hetSte* relative to *euSte* (17% vs. 4%). Intriguingly, *euSte* exhibits a bimodal copy number distribution, with the majority of lines carrying fewer than 25 copies but with 17% carrying more than 150 (Fig. 2). We reasoned that the two peaks of this distribution correspond to the classical *Ste* (high copy number) and *Ste+* (low copy number) alleles previously described^13^ on the basis of Ste crystal morphology in the testes of males lacking *Su(Ste)*, as the frequencies we observed for these copy-number alleles are strikingly similar to those reported in Palumbo et al. (1994). Copy numbers of *Su(Ste)* and *hetSte*, on the other hand, are unimodal and appear to be approximately normally distributed (Fig. 2). There is also substantial interpopulation differentiation with respect to copy number of these elements. High copy number *euSte* (>100 copies) alleles are common in four out of the five GDL populations but are absent in the Zimbabwe lines sampled, all of which carry 10 or fewer copies or fewer. Interestingly, *hetSte* is also less abundant in the Zimbabwe population as compared to all others (*t* = 2.75; *p* = 0.0042). *Su(Ste)* also exhibits clear patterns of population differentiation, with Ithaca and Tasmania lines carrying higher copy numbers than the remaining populations and Beijing lines carrying the fewest copies (Fig. 2). Furthermore, *Su(Ste)* exhibits substantial variation in sequence, forming three distinct clades (also see Chang & Larracuente (2019); Supplemental Fig. 2). Interestingly, one of these clades (clade 3 in Supplemental Fig. 2) is underrepresented in the Beijing population relative to all others (*t* = 3.24284; *p* = 0.0017; Supplemental Fig. 3).

**Figure 2:**
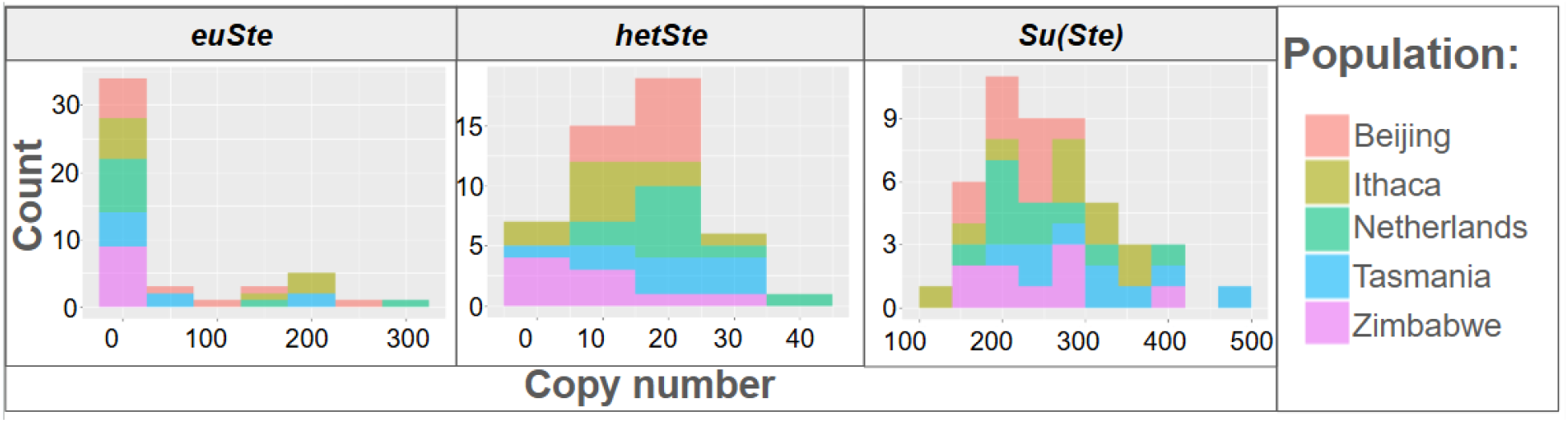
Histograms of copy number for *Ste-Su(Ste)* family elements across a panel of 48 Global Diversity Lines estimated by calculating read depth to element consensus sequences for each element, relative to a panel of single-copy genes on the same chromosome. Histograms from left to right: *euSte, hetSte, Su(Ste)*. Colors denote population of origin.

### Males carrying high *Stellate* copy number X chromosomes produce biased sex ratios

The observed patterns of copy number variation led us to hypothesize that *euSte* may be an active driver in contemporary populations, with high copy number alleles potentially overcoming host suppression in a dose-dependent manner. To test this hypothesis, we generated F1 males from 56 pairwise combinations of GDLs involving 4 naturally-occurring high *euSte* copy number X chromosomes and 14 low *Su(Ste)* copy number Y chromosomes (Fig. 3A) at three temperatures (18, 25, and 29°C). These F1 males were subsequently screened for X chromosome drive by scoring the sex ratio in the F2 generation as a proxy for the relative transmission of the X and Y chromosomes (though see discussion of sex chromosome nondisjunction below). A substantial overrepresentation of daughters was observed in a subset of genotypes at the lowest temperature tested (18°C; Fig. 3B). Strikingly, this was primarily (though not exclusively) observed in F1 males carrying Y chromosomes from the Beijing population. In most genotypes, the distortion observed was moderate (<60% females), with the strongest sex-ratio skew observed being 65% females. On the other hand, at 25°C and 29°C, the vast majority of genotypes exhibited approximately Mendelian sex ratios, though at 29°C the entire distribution is slightly skewed toward excess females (Fig. 3B). Taken together, these crosses strongly suggest that *Ste* arrays can cause sex-ratio distortion in crosses between contemporary strains. (For all individual crosses, associated *p*-values obtained from an Exact Binomial test for deviation from a 50-50 sex ratio and subject to FDR correction can be found in Supplemental File 1, Table 2.)

**Figure 3:**
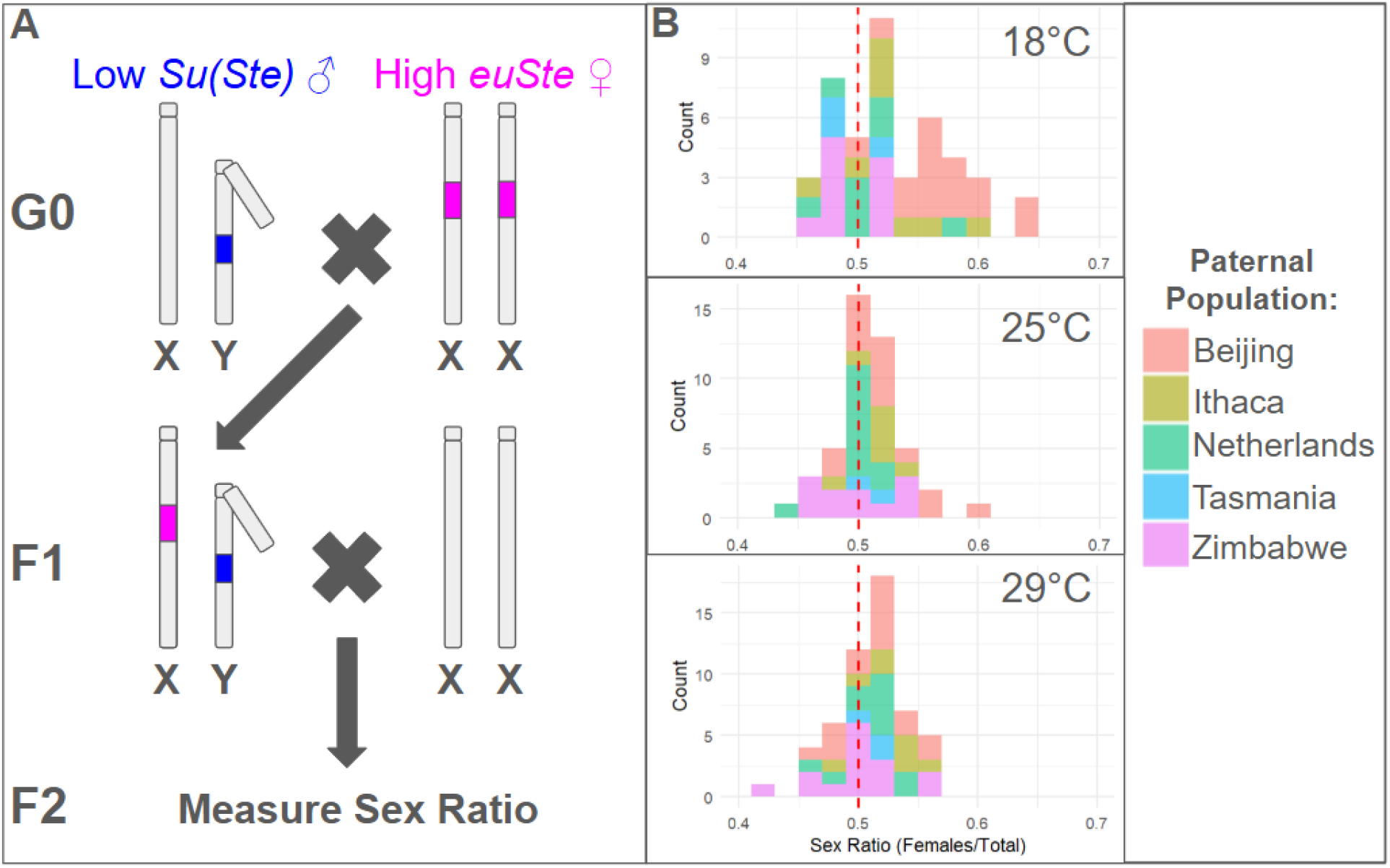
Crosses to screen for evidence of sex ratio distortion occurring in F1 males obtained via intercrossing of GDLs. A) High *euSte* copy-number females (4 lines) were crossed to low *Su(Ste)* copy-number males (14 lines) to generate F1 males with high *Ste* and low *Su(Ste)* copy number. These males were subsequently outcrossed to standard reference line females and screened for evidence of drive through assessment of the F2 sex ratio. B) Histograms of sex ratio produced by F1 males at 3 temperatures. Colors represent the population of origin of the G0 male parent and indicate a pronounced effect of paternal background (i.e. the source of the Y chromosome in the F1 males screened for drive) in predicting sex ratio.

### Deletion of the *euSte* array restores Mendelian inheritance of sex chromosomes

To definitively prove that the observed sex ratios are *Ste*-dependent, we deleted the entire *euSte* array in the high *Ste* copy number line B11 (240 copies; ~275 Mb) and replaced it with a fluorescent transgene via CRISPR/Cas9-mediated homologous recombination (Fig. 4A). The resulting line is genetically identical to B11 aside from this deletion, allowing us to isolate the effect of the presence/absence of the *euSte* array from all other variables. This is especially important given that many backgrounds with high *Ste* copy number exhibit no drive. We repeated a subset of crosses showing skewed sex ratios using B11 with and without the *euSte* array and confirmed that, in all cases, Mendelian segregation was restored by the deletion. This indicates that a high copy number of *euSte* is necessary for the observed sex-ratio distortion phenotype (Fig. 4B).

**Figure 4:**
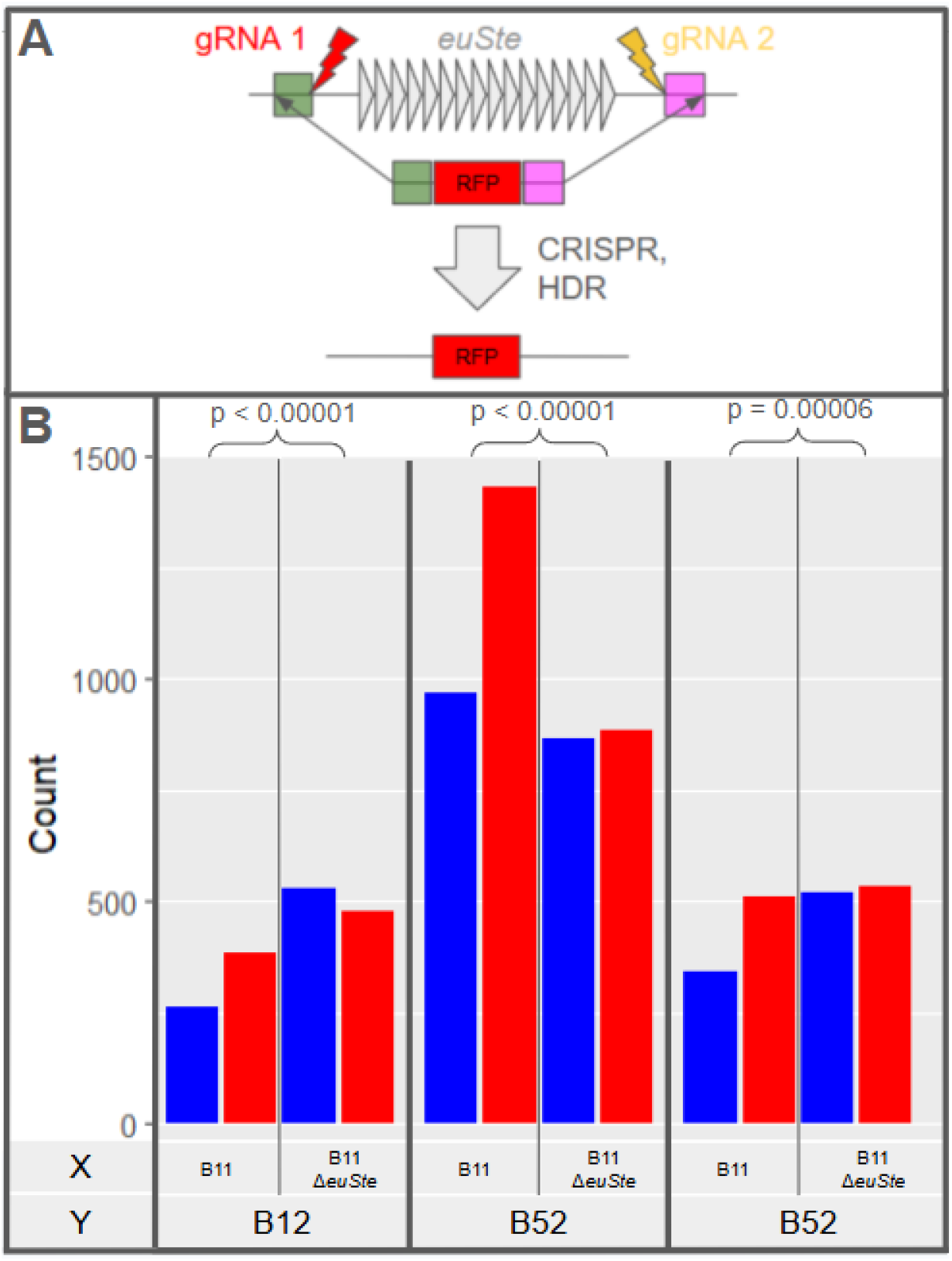
Rescue of drive via precise deletion of the *euSte* array in an otherwise matched genetic background. A) A naturally-occurring high copy number *euSte* allele was deleted in a single step via CRISPR-Cas9 by cutting unique sites up- and downstream of the array followed by HDR-mediated integration of a *MHC-dsRed* fluorescent transgene in its place. B) counts of male (blue) and female (red) offspring obtained from F1 males carrying either the unedited driving *euSte* allele or the matched deletion across three genetic backgrounds where we had previously identified drive. In all cases, the aberrant sex ratio is rescued in the absence of *euSte*. p-values were obtained from a chi square test performed on a 2×2 contingency table of counts of male and female offspring.

### *euSte* expression level does not predict drive

To investigate the source of variation among genotypes in sensitivity to *euSte-*dependent drive, we performed RT-qPCR to measure the abundance of spliced *euSte* mRNAs in a subset of genotypes. Surprisingly, although *euSte* copy number predicts expression level as expected, expression level shows no meaningful correlation with the degree of meiotic distortion (Fig. 5).Rather, F1 males carrying the same *euSte* allele exhibit very similar *Ste* expression levels regardless of whether they produce biased sex ratios.

**Figure 5:**
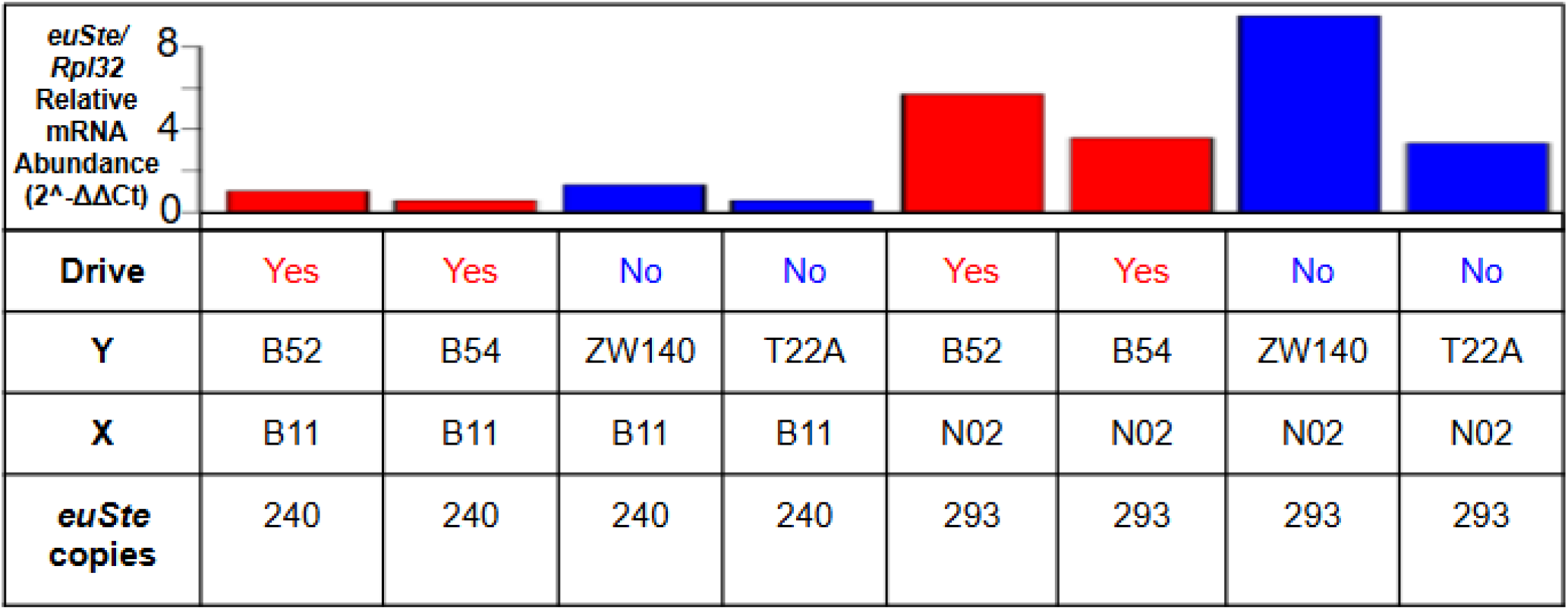
The abundance of spliced *euSte* mRNA in testes across genotypes with (red) or without (blue) sex ratio meiotic drive assessed via RT-qPCR and quantified relative to the ribosomal protein gene *Rpl32*. For the ease of interpretation of relative expression across lines, we arbitrarily set the *euSte* expression level observed in B52/B11 F1 males (column 1) to a value of 1 and adjusted the values for the other lines proportionally. The 8 F1 male genotypes tested arise from all pairwise combinations of two high *euSte* copy number X chromosomes and four Y chromosomes (as well as the accompanying genetic backgrounds), two sensitive to drive and two insensitive.

## Discussion

Having established that *Ste* is an active driver in contemporary populations, the patterns of copy number variation we observed for *Ste*-family elements raise intriguing questions about the evolutionary history of the system, the pressures that maintain its current patterns of polymorphism, and the functional consequences of natural variation in its components. While population genetic theory and associated modelling support the idea that meiotic drive conflicts are expected to be short and intense^e.g. 18,35^, the intermediate frequency of high copy number *euSte* alleles observed across most populations instead suggests that the system may currently exist at a stable equilibrium. This pattern is reminiscent of the other known driver in *D. melanogaster, Segregation distorter* (*SD*). Although the frequency of SD is lower (~1–5%), SD allele frequencies are also largely stable over time and across populations^36^. The most widely accepted explanation for the apparent stable equilibrium of SD is that it arises from a trade-off between drive and its concomitant costs to fertility and viability. While *Ste’s* effects on fertility and sperm competitive ability in wildtype animals are unexplored, the fertility costs associated with derepression of *Ste* in mutant backgrounds deficient in *Su(Ste)* or lacking components of the piRNA pathway are well-documented^5,19^. Therefore, a similar mechanism may be at play. The absence of high copy number alleles in the Zimbabwe population is unexpected and may simply reflect the relatively small number of lines sampled. On the other hand, it is also possible that the frequency of driving *Ste* alleles varies with latitude, possibly due to the temperature dependence of *Ste* drive, as observed with other X chromosome drivers in other *Drosophila* species^37^. While further study is needed, reanalysis of an existing (albeit geographically limited) allele-frequency dataset^19^ suggests the existence of such a cline, with the frequency of high copy number (classical *Ste)* alleles increasing with distance from the equator (Supplemental File 1, Table 6; *p* = 0.046; Spearman’s rho = 0.6105).

While we have demonstrated that high *euSte* copy number is necessary for drive in natural backgrounds, the role of *hetSte* in the system is less clear. Given its almost 10-fold lower maximum copy number, the higher incidence of degenerate copies relative to *euSte*, and its unimodal CNV distribution, we hypothesize that this array may represent an earlier invasion of *Ste*-family elements that subsequently lost the ability to drive (at least autonomously) due to escalating host suppressive capacity, necessitating either an increase in *Ste* dosage or a change in protein sequence for reestablishment of biased segregation. Innovations in these features may have allowed for *euSte’s* subsequent invasion, though other models can be imagined (a complication to this particular hypothesis is the presence of a *Ste*-like pseudogene near the syntenic location of the *euSte* array in *D. simulans*^24^, indicating that *euSte* may actually be the older of the two elements). In any case, because X-linked sequences like *hetSte* benefit from *euSte* drive and are thus expected to cooperate with driving *euSte* alleles, the intact coding potential of most *hetSte* copies in our dataset combined with earlier observations^38–39^ of derepression of these copies in permissive mutant backgrounds suggest the possibility that *hetSte* may still positively modify driving *euSte* alleles. It could function in this capacity either by simply contributing additional *Ste* protein (if hetSte protein is capable of producing drive) or through the production of additional *Ste* mRNAs, which could act as a “sponge” to soak up *Su(Ste)* piRNAs, boosting the expression of driving *euSte* alleles. Indirect support for shared selective pressures between *euSte* and *hetSte* (as would be expected for a cooperative relationship) comes from the fact that in the Zimbabwe population, from which we recovered no high copy number *euSte* alleles, *hetSte* copy number is also significantly lower than the other populations.

It is notable that, while most known drivers produce strong distortion, the sex ratio distortions observed here were generally less than 60% and highly dependent on both temperature and genetic background. These features may explain why *Ste* was long incorrectly assumed to be inactive despite extensive study, an easily scorable phenotype by which to detect drive (sex ratio), and frequent high copy number alleles in natural populations and standard lab stocks alike. The fact that such a system could be overlooked in even *D. melanogaster* suggests that other weak drivers may remain to be discovered even in the best studied models. It also raises the possibility that other “cryptic” drive systems, assumed to be wholly inactive due to the presence of host suppression, may actually exist in a state similar to *Ste*, where weak drive may still actively shape the evolution of the system itself and the host genome at large despite effective suppression in most contexts. *But why would host suppression remain incomplete?* One possibility is that pleiotropic costs of perfect suppression outweigh the costs of weak drive in some situations. For example, perhaps complete *Ste* suppression via extensive *Su(Ste)* piRNA production might reduce the availability of the effectors of the piRNA pathway, Aub and AGO3, to effectively suppress other targets, like transposons. Indeed, upregulation of these factors in *D. melanogaster* is necessary to prevent *Ste*-associated sterility^5^, suggesting that they must have been rate-limiting at some points in the conflict. Additionally, modelling suggests that a female-skewed sex ratio may actually be adaptive under some circumstances, meaning that alleles permissive to weak drive may sometimes be selectively favored (so long as they are not Y-linked)^40–41^.

The observation that the female:male sex ratio does not exceed 2:1 is consistent with the mechanism of *Ste* drive proposed by Meng & Yamashita (2025; outlined in Fig. 1B), which predicts precisely this ceiling for drive strength (although note that more extreme ratios have been previously reported from *Su(Ste)* mutants^19^). Interestingly, most genotypes tested here did not approach this theoretical limit. Among the normally disjunct products of a single primary spermatocyte, the only X:Y transmission ratios possible are 1:1 (Mendelian), 2:1 (if Ste protein associates with the Y-bearing cell in meiosis I and segregates asymmetrically in meiosis II), or 1:0 (if Ste segregates with all Y-bearing spermatids and kills them). This implies three possible mechanisms for intermediate ratios to occur (i.e. between 1:1 and 2:1): 1) variation in the abundance of Ste protein or of factors that modify sensitivity to its toxicity among spermatids, primary spermatocytes, or cysts; 2) variation in the cosegregation of Ste protein with the Y chromosome among meioses; or 3) that some or all Ste-bearing sperm are viable and fertilize at an appreciable rate but suffer a sperm competitive disadvantage relative to their non-Ste-bearing counterparts. While these scenarios are not mutually exclusive, given the extreme cytological defects reported by Meng & Yamashita (2025) in most Ste-bearing spermatids, we favor the former two possibilities (or some combination thereof). It is also possible that this effect is explained at least in part by the elevated rate of XY nondisjunction observed upon derepression of *Ste*. In these cases, drive overwhelmingly (>10:1) favors nullo-over XY-bearing sperm, meaning that the vast majority of nondisjunct progeny are male, thus reversing the sex ratio effect. This observation also suggests that Ste’s asymmetric segregation and/or toxicity may be modulated by the differential in total chromatin, or in the abundance of certain sequence features, between reciprocal products. The rate of nondisjunction due to *Ste* has been quantified only in *Su(Ste)* mutants^19^, where it may be much higher than in wildtype flies undergoing *Ste* drive. Our scheme, however, does not allow differentiation of nondisjunct classes, so the rate of nondisjunction in natural backgrounds and its contribution to the intermediate sex ratios observed cannot be inferred.

The genetic determinants of the observed variation in drive among genotypes and the mechanisms by which they modify the penetrance of *Ste* remain unknown. We propose several non-exclusive contributors: 1) Y-linked variation in *Su(Ste)* copy number and sequence; 2) variation in whatever broad features of the Y chromosome mediate Ste protein’s preferential cosegregation with it; 3) variation in components of the piRNA pathway and in the maternally deposited piRNA pool; and 4) variation in the expression and/or sequence of genes that modify the segregation or toxicity of Ste protein. We find that the paternal background explains much of the variation observed, suggesting a pronounced Y-chromsome effect (Supplemental File 1, Table 4). However, contrary to our expectation, we observed no correlation between *euSte* mRNA abundance with the strength or presence/absence of drive. Whether or not this measure corresponds meaningfully to Ste protein abundance is unknown, but in either case it suggests the existence of additional Y-linked modifiers that act downstream of modulation of *Ste* mRNA abundance by *Su(Ste)* piRNAs. Given the paucity of Y-linked protein coding genes, we propose that the extensive repetitive variation among Y chromosomes in natural populations may affect the ability of *Ste* protein to target and/or damage Y-bearing sperm.

## Materials and Methods

### Estimation of the copy number of *Ste*-family elements in the GDL

For copy number estimation across the GDL, we made use of PacBio sequence data representing 48 lines as well as associated assemblies of 27 of these lines (see Data Availability for information about deposition of these datasets). We first estimated the copy number of these elements from the raw sequencing reads via read depth. DNA consensus sequences for *euSte, hetSte, Su(Ste)*, and *pseudo-CK2βtes-Y* (*PCKR*; a paralogous family of Y-linked pseudogenes^20^) were generated as follows; a 0.43 kb region of shared homology between all *Ste*-family elements was obtained from all copies identified from the assemblies as described above. The resultant sequences were aligned using MAFFT^42^ and a consensus inferred using Jalview^43^. Sites containing >50% gaps were removed. The reads were subsequently aligned using minimap2 (parameters: -t 6 -B 2 -ax map-hifi) to a FASTA file containing these consensus sequences as well as fragments of similar size derived from exons of X- and Y-linked single-copy coding genes for normalization (Supplemental File 2).

For the subset of sequenced lines with highly complete assemblies, copy number estimation of *Ste*-family elements was also carried out via blast and phylogenetic clustering of hits. A 0.43 kb section of the *euSte* coding sequence with homology to all *Ste*-family elements was obtained from FlyBase. A blastn^44^ search using this query identified similar sequences in the assemblies described above. Hits from all assemblies were subsequently aligned with MAFFT^42^. An approximately maximum likelihood phylogeny was constructed from this alignment using FastTree^45^. Clades corresponding to *euSte, hetSte, Su(Ste)*, and *pseudo-CK2β* repeats (PCKR; paralogous Y-linked pseudogenes) were manually annotated in figtree and the sequences corresponding to each clade recovered using TREE2FASTA^46^.

### Fly crosses to screen for evidence of meiotic drive

See Figure 2A for an overview of the mating scheme. Stocks were maintained at 25°C prior to starting the experiment. For each temperature condition (18, 25, and 29°C), both the G0 and F1 crosses were maintained at these temperatures for their entire life cycles. G0 crosses contained 5 males and females, flipped weekly, while F1 crosses contained 5 males and females and were flipped twice weekly and discarded after the fifth flip. After removing the parents, the F2 flies were moved to 25°C, and scored for sex. Vials were discarded prior to the eclosion of the F3 generation. For each genotype-temperature combination (*n* = 168), counts were only included in the final dataset if at least 150 adults were scored (*n* = 144; mean adults counted per condition = 540).

### Deletion of *euSte* array in GDL B11

We designed and synthesized gRNA and homology arm dsDNA constructs (Twist Biosciences; Supplemental File 1, Table 8) and subsequently integrated them into pJAT30 (Addgene plasmid # 204289), a kind gift from David Stern, via double Gateway cloning using Gateway LR Clonase (Invitrogen)^47^. The resultant plasmid contains sequences driving expression of two gRNA targeting sites flanking the *euSte* array as well as a homology directed repair donor carrying an *MHC-dsRed* transgene flanked by homology arms matching the sequences up and downstream of the intended breaks (Fig. 4A). This plasmid was subsequently coinjected alongside a plasmid containing a *nos-Cas9* transgene (all embryo injections were performed by Rainbow Transgenic Flies Inc.) into GDL B11, which carries 240 *euSte* copies in tandem. We identified putatively *euSte-*deleted transformants via red fluorescence and subsequently confirmed the deletion via PCR (Supplemental Fig. 1; primers in Supplemental File 1, Table 7).

### Rescue experiments

For a subset of genotypes exhibiting drive, we repeated the crosses as described above at 18°C in parallel using both GDL B11 and the background-matched *euSte* deletion line as the source of the X chromosome in the F1 males to be screened for drive.

### Quantification of *euSte* mRNA

We constructed F1s representing 8 pairwise combinations of two high *euSte* copy number X chromosomes (from B11 and N02, with 240 and 293 copies respectively) and four Y chromosomes, of which two are sensitive to drive (B52, B12) and two are resistant (ZW140, T22A). Flies were reared at 18°C. To assess *euSte* mRNA abundance, we extracted RNA from the dissected testes of 15 males per genotype using QIAzol (QIAGEN) according to manufacturer instructions. cDNA was subsequently synthesized using Protoscript II Reverse Transcriptase (New England Biosciences) with d(T)23VN primers to selectively amplify polyadenylated transcripts. RT-qPCR was carried out using LightCycler® 480 SYBR Green I (Roche) to quantify the abundance of *euSte* relative to the housekeeping gene Rpl32 (primer sequences can be found in Supplemental File 1, Table 7^29,32^).

### Reanalysis of an existing allele frequency dataset

Using the frequency data for *Ste* and *Ste+* alleles reported in Palumbo et al. (1994) from various localities (primarily in Italy), we tested via Spearman’s rank correlation whether there was a correlation between latitude and *Ste* frequency.

### Statistical analysis

Statistics were carried out using Rstudio (Supplemental File 4). To assess the concordance of copy number estimates derived from different methods (read depth vs. blast and phylogenetic clustering), we applied spearman’s rank correlation. In the cross experiment to screen for evidence of drive, we tested for significant deviations from a 50:50 sex ratio via exact binomial test with FDR correction applied. To assess the role of temperature as well as maternal and paternal background in predicting sex ratio in the same dataset, we applied a binomial GLM. To assess rescue of drive upon deletion of the *euSte* array, we performed a chi square test on a 2×2 contingency table of counts of male and female offspring from pairs of matched genotypes with and without *euSte*. For testing whether *hetSte* is less abundant in Zimbabwe than all other populations, we used a one-tailed two sample t-test.

## Data availability statement

All code and data required to reproduce the analyses and figures presented here can be found in the supplemental files. The associated sequencing reads and assemblies will be deposited prior to publication (but are not yet deposited at the time of submission).

---

**Supplemental Figure 1:**
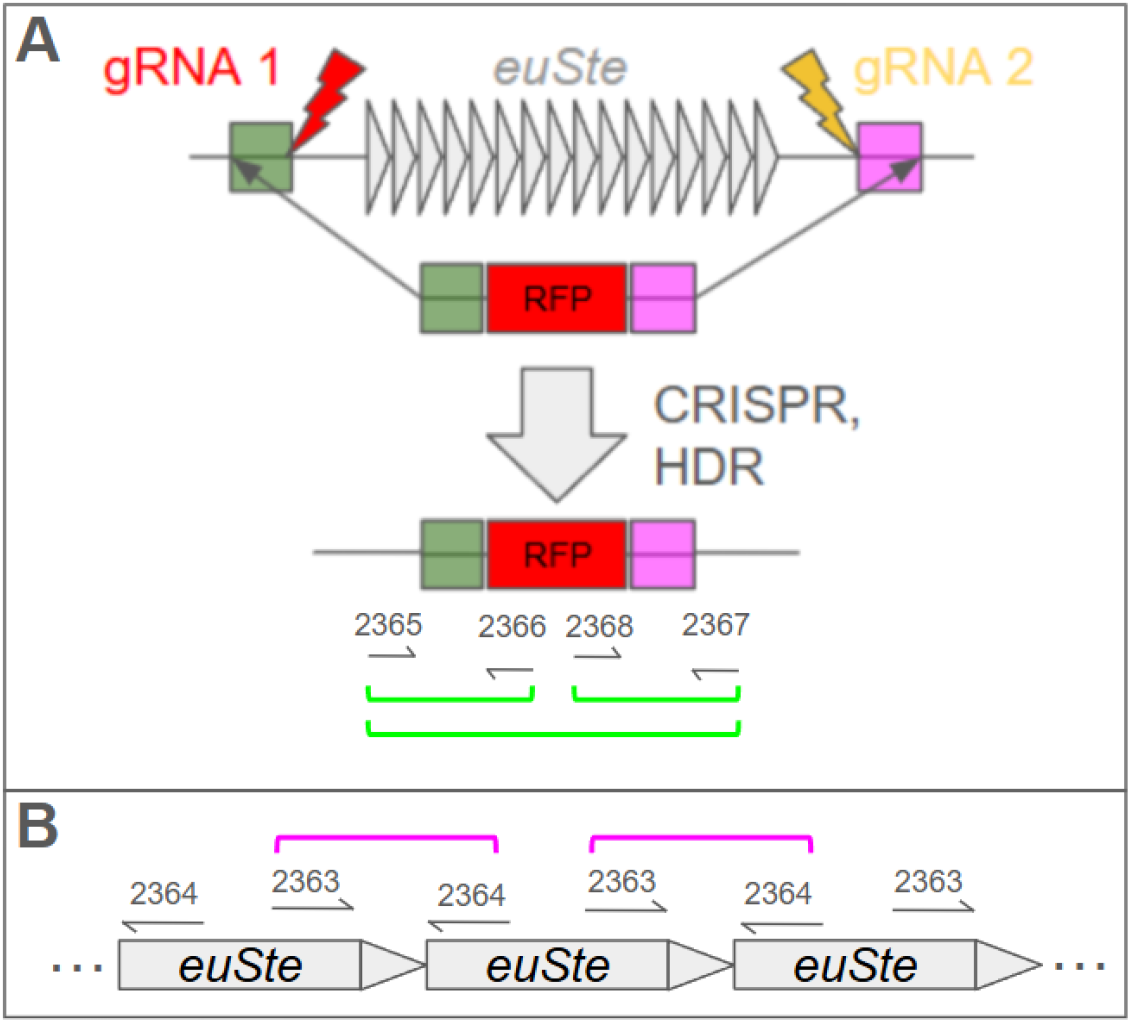
Primers used to confirm deletion of the *euSte* array in GDL B11. Brackets represent primer pairs tested and the resulting amplicons; green brackets denote pairs that amplify from the deletion line but not B11, while pink brackets indicate pairs that amplify from B11 but not the deletion line. A) Primers used to confirm proper integration of transgene; B) Primers used to confirm absence of tandem *euSte* copies. Note that only 3 of 240 copies are shown.

**Supplemental Figure 2:**
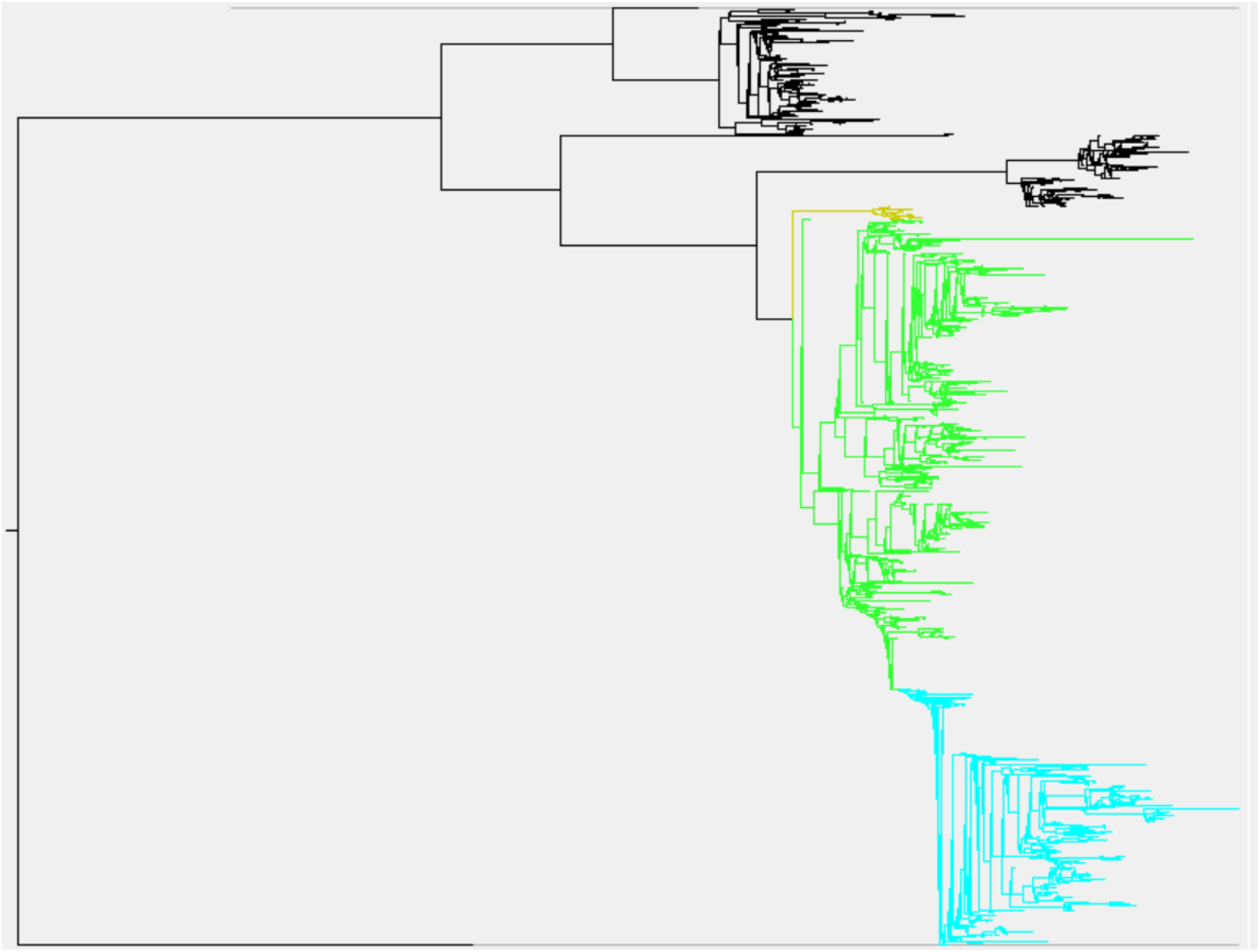
Approximately maximum likelihood tree (FastTree) of *Ste*-family elements (*PCKR, euSte, hetSte, Su(Ste)*; outgroup is *SSL*, the closest single-copy autosomal paralog to these genes) from 27 GDL lines with the three major clades of *Su(Ste)* shown in yellow (clade 1), green (clade 2), and cyan (clade 3). The corresponding tree file can be found in Supplemental File 7.

**Supplemental Fig. 3:**
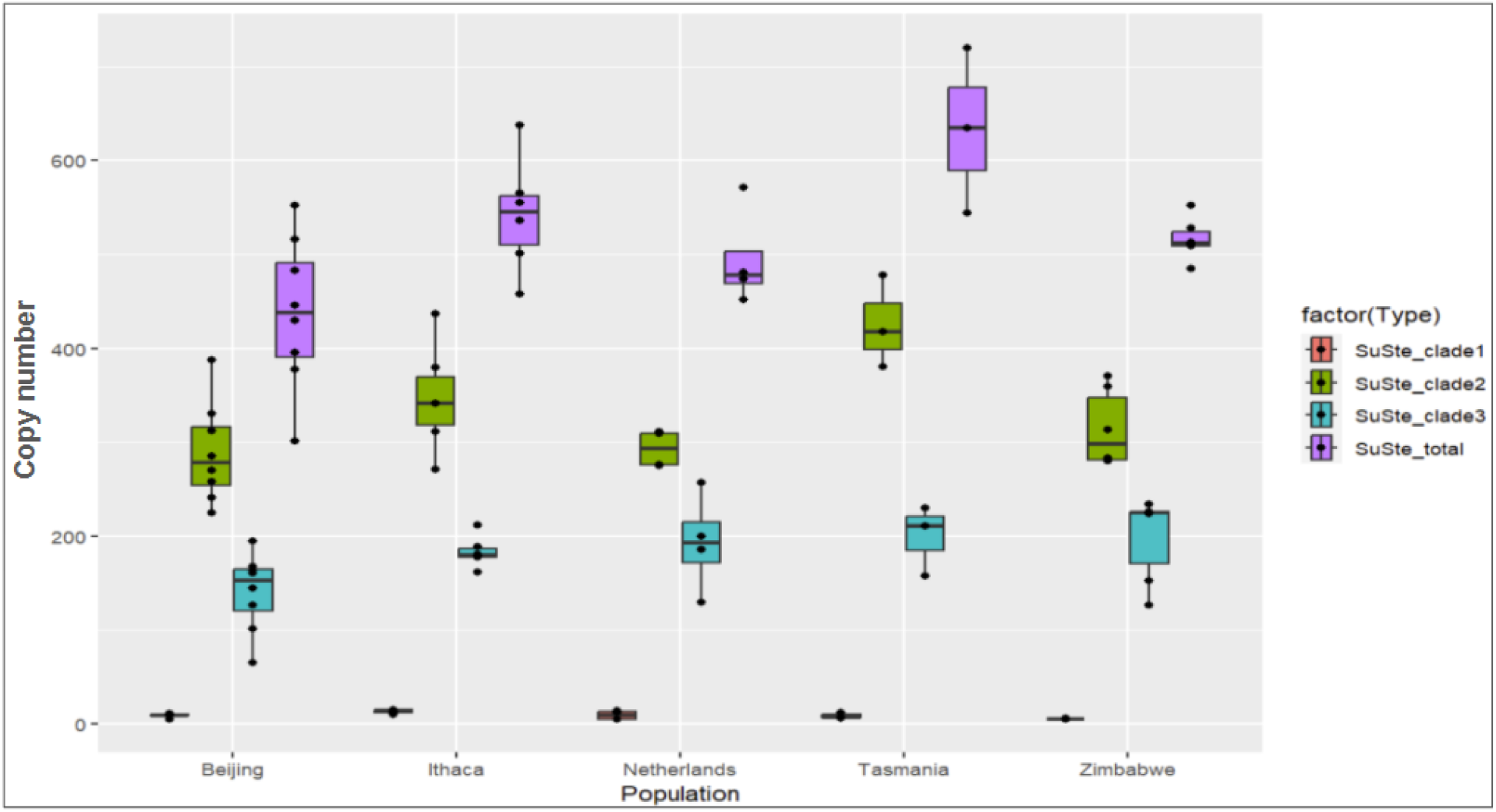
Copy number estimates for *Su(Ste)* (both total and by clade) obtained via blast and phylogenetic clustering of copies identified from assemblies. See Supplemental Fig. 2 for the relationship among the clades.

## Supplemental Files

- Supplemental File 1: Supplemental Tables 1-11
- Supplemental File 2: Fasta containing consensus sequences used in read depth-based copy number estimation
- Supplemental File 3: Blast query used in assembly based copy number estimation
- Supplemental File 4: R code underlying plots and statistics
- Supplemental File 5: Bash code for estimating copy number from read depth
- Supplemental File 6: Bash code for estimating copy number from assemblies

