## Supplemental File 6 for "The *Stellate* meiotic drive system of *Drosophila melanogaster* is active in contemporary populations"

###COPY NUMBER ESTIMATION FROM ASSEMBLIES
ls *hifi*.fasta | parallel -j 10 'blastn  -query 01_Ste_homology_only_query.fasta  -subject {} -word_size 11 -evalue 0.05 -gapopen 5 -gapextend 2 -reward 2 -penalty -3 -outfmt 6 -out {.}.homology.only.blastn.txt' &
#This copies the filename to the beginning of each line for tracking of which sequences arose from which assembly in subsequent trees/alignments
for filename in ./*hifi*.txt
do
    grep -H ^ *hifi*.txt > output.txt
done;
#Replaces filename in table with original genome filename; adds tab
sed -i 's/.homology.only.blastn.txt:/.fasta	/g' output.txt 

#fix line break encoding (dos vs. unix)
for filename in ./*hifi*.bed
do
	tr -d '\015' <$filename >"${filename}.fixed"
done;

#Separate forward and reverse hits
for filename in ./*hifi*.bed
do
   awk '{OFS="\t"; if ($3-$2 > 0) {print}}' $f >"${filename}_FORWARD.bed"
   awk '{OFS="\t"; if ($2-$3 > 0) {print}}' $f >"${filename}_REVERSE.bed"
done;

#Switch columns 2 and 3 for reverse hits so that all end coordinates are greater in value than the start 
for filename in ./*hifi*REVERSE.bed.fixed
do
    awk ' { OFS="\t"; t = $3; $3 = $2; $2 = t; print; } ' $filename >"${filename}.rev"
done;

#manually filter out any hits w/ <60% query cover in excel
#Once you've taken the output file, thresholded QC in excel, removed all columns but 'genome_file' 'chrom' 'chromStart' 'chromEnd', separated back out by genome in R and exported as BED file,
#This code will output hits to fasta file
ls *.fasta | cut -f 1 -d . | parallel -j 10  'bedtools getfasta -fo {.}_FORWARD.fasta -fi {}.fasta -bed {}_FORWARD.bed.fixed' &
ls *.fasta | cut -f 1 -d . | parallel -j 10  'bedtools getfasta -fo {.}_REVERSE.fasta -fi {}.fasta -bed {}_REVERSE.bed.fixed.rev' &

#Reverse complement reverse fasta files
export PATH=/programs/seqtk:$PATH
for filename in ./*REVERSE.fasta
do 
	seqtk seq -r $filename >"${filename}.comp"
done;

#Add genome file name to fasta headers (for tracking of which sequence came from which assembly)
for f in *FORWARD.fasta
do 
	sed -i "s/^>/>${f}_/" "$f"
done;
for f in *REVERSE.fasta.comp
do 
	sed -i "s/^>/>${f}_/" "$f"
done;

#Combine all forward fastas, then all reverse, then combine these outputs
cat *FORWARD.fasta > all_F.fasta
cat *REVERSE.fasta.comp > all_R.fasta
cat all*.fasta > all.fasta

#Align with MAFFT
export PATH=/programs/mafft/bin:$PATH
mafft --thread 20 all.fasta > all_aligned.fasta

#Fixing sequence names in alignment to pass through fasttree (remove ':')
sed -i 's/:/_/g' all_aligned.fasta 

#make fasttree tree
export PATH=/programs/FastTree-2.1.11:$PATH
FastTree -gtr -nt < all_aligned.fasta > all_tree

#Make tree, color taxa in figtree by type (e.g. Su(Ste)); run TREE2FASTA locally (NOT ON lm06) to separate out clades
perl TREE2FASTA_v1_6.pl all_tree_colored.nex all_aligned.fasta

#Name each file list_[element_name] (e.g. list_euSte)

#name of each sequence in tree is a bed file with the columns separated by extra underscores; this step turns the sequence names back into bed files of their locations
for f in list*
do 
	sed 's/.fasta_/.fasta	/' $f >"${f}_fixed"
done;

for f in list*fixed
do 
	sed 's/.fasta./.fasta	/' $f >"${f}_again"
done;

for f in list*again
do 
	sed 's/comp_//' $f >"${f}_twice"
done;

for f in list*twice
do 
	rev "$f" | sed 's/_/\t/' | rev >"${f}_table"
done;

for f in list*table
do 
	rev "$f" | sed 's/-/\t/' | rev >"${f}.bed"
done;

#making table of counts (homology only)
#step1 make list of GDL lines (contained in file line_list.txt
ls | cut -f 1 -d _ > line_list.txt
#step2 make list of counts for each element type (NOTE: for lines with 0 copies of a given element, no number is outputted)
fgrep -of line_list.txt list_euSte_fixed_again_table.bed | sort | uniq -c | awk '{print $1}' >euSte_counts.txt
fgrep -of line_list.txt list_hetSte_fixed_again_table.bed | sort | uniq -c | awk '{print $1}' >hetSte_counts.txt
fgrep -of line_list.txt list_SuSte_fixed_again_table.bed | sort | uniq -c | awk '{print $1}' >SuSte_counts.txt
