## Supplemental File 5 for "The *Stellate* meiotic drive system of *Drosophila melanogaster* is active in contemporary populations"

###COPY NUMBER ESTIMATION FROM READ DEPTH TO CONSENSUS SEQUENCES
#map reads to consensus sequences
export PATH=/programs/minimap2-2.26:$PATH
export PATH=/programs/samtools-1.9-r9/bin/samtools:$PATH
nohup sh -c 'ls *fastq.gz | parallel -j 2 "minimap2 -t 6 -B 2 -ax map-hifi --sam-hit-only Consensus_Sequences_homology_v1.fasta {} | samtools sort -@ 6 -o {.}_sort.bam"' &
nohup sh -c 'ls *sort.bam | parallel -j 6 "samtools index -b -o {.}.bai {}"' &
export LC_ALL=en_US.utf-8
export LANG=en_US.utf-8
export PATH=/programs/deeptools-3.5.1/bin:$PATH
export PYTHONPATH=/programs/deeptools-3.5.1/lib64/python3.9/site-packages:/programs/deeptools-3.5.1/lib/python3.9/site-packages
ls *.bam | parallel -j 6 'bamCoverage --bam {} -o {.}_coverage_full.bedgraph --numberOfProcessors "6" --binSize "650" --normalizeUsing "CPM" --outFileFormat "bedgraph"' &
ls *.bam | parallel -j 6 'bamCoverage --bam {} -o {.}_coverage_bin.bedgraph --numberOfProcessors "6" --binSize "10" --normalizeUsing "CPM" --outFileFormat "bedgraph"' &

#Analyzing data for full.bedgraph files (a single bin per consensus)
#get only fourth column with read depth (CPM) for, in order, SuSte (1), PCKR (2), hetSte (3), euSte (4), Sxl (5), white (6), yellow (7), FDY (8), ARY (9), PRY (10)
for f in ./*full.bedgraph
do
	awk '{ print $4 }' $f >"${f}_depth"
done; 

#replace line breaks with tab
for f in ./*full.bedgraph_depth
do
	awk -vRS="\n" -vORS="\t" '1' $f >"${f}_transformed"
done;

#get read depth normalized to either ARY + PRY (SuSte/PCKR) or Sxl + w + yellow (euSte, hetSte); output is normalized count for in order: SuSte, PCKR, hetSte, euSte
for f in ./*full*transformed
do
	awk '{OFS="\t"; print $1/($9/2 + $10/2), $2/($9/2 + $10/2), $3/($5/3 + $6/3 + $7/3), $4/($5/3 + $6/3 + $7/3)}' $f >"${f}_counts"
done;

#replace tabs with line breaks
for f in ./*full*counts
do
	sed 's/\t\t*/\n/g' $f >"${f}_turned.txt"
done;

#combine into one file and add filename
for filename in ./*turned.txt
do
    grep -H ^ *turned.txt > output.txt
done;

#Delete part of the filename appended to the beginning of each line in the last code chunk to separate columns
sed -i 's/_sort_coverage_full.bedgraph_depth_transformed_counts_turned.txt:/	/g' output.txt 
